# Polyclonal epitope cartography reveals the temporal dynamics and diversity of human antibody responses to H5N1 vaccination

**DOI:** 10.1101/2020.06.16.155754

**Authors:** Julianna Han, Aaron J. Schmitz, Sara T. Richey, Ya-Nan Dai, Hannah L. Turner, Bassem M. Mohammed, Daved H. Fremont, Ali H. Ellebedy, Andrew B. Ward

## Abstract

Novel influenza A virus (IAV) strains elicit recall immune responses to conserved epitopes, making them favorable antigenic choices for universal influenza virus vaccines. Evaluating these immunogens requires a thorough understanding of the antigenic sites targeted by the polyclonal antibody (pAb) response, which single particle electron microscopy (EM) can sensitively detect. Here, we employed EM polyclonal epitope mapping (EMPEM) to extensively characterize the pAb response to hemagglutinin (HA) after H5N1 immunization in humans. Cross-reactive pAbs originating from memory B cells immediately bound the stem of HA and persisted for over a year post vaccination. In contrast, *de novo* pAb responses to multiple sites on the head of HA, which targeted previously determined key neutralizing sites on H5 HA, expanded after the second immunization and waned quickly. Thus, EMPEM provides a robust tool for comprehensively tracking the specificity and durability of immune responses elicited by novel universal influenza vaccine candidates.

## INTRODUCTION

Despite decades of endeavors to produce a lasting therapeutic and effective vaccine, seasonal influenza virus still causes a tremendous burden to public health each year, and pandemic influenza virus is a constantly looming threat. To understand the range of protection needed for seasonal and universal influenza virus vaccines (vaccines that can generate protection against a broad array of influenza virus strains), we need a thorough characterization of humoral immune responses to influenza virus vaccination (Erbelding *et al.*, 2018).

Because of antigenic drift, the ability of influenza virus to mutate in response to selection pressures imposed by host immune systems, seasonal influenza virus vaccines must be reformulated yearly and still result in only 10-60% efficacy (Centers for Disease Control and Prevention, 2018). Current vaccines are strain-specific, eliciting antibody responses primarily to the variable head region of hemagglutinin (HA). This creates a challenge for generating vaccines to potentially pandemic strains, since it is almost impossible to predict which strain can cause a pandemic.

Highly pathogenic avian influenza (HPAI) H5N1 viruses have been periodically crossing the species barrier from birds into humans, causing serious lower respiratory tract infections and viral pneumonia (Claas *et al.*, 1998; Peiris, De Jong and Guan, 2007; Cowling *et al.*, 2013). As the human population is largely naïve to these avian IAV strains, humans have little pre-existing immunity to H5N1 infections, resulting in a devastating 50-60% mortality rate (WHO, 2020). However, due to frequent exposure to seasonal influenza virus HAs, humans harbor memory B cells that are directed against epitopes shared between such HAs and H5 HA. These epitopes predominantly reside within the conserved stem region of HA, and are the targets of broadly reactive antibodies (Throsby *et al.*, 2008; Ekiert *et al.*, 2009). A major strategy for universal influenza vaccine design is to re-focus immune responses to the immuno-subdominant but conserved stem by immunizing with HA from non-circulating influenza virus strains, such as H5N1, thereby recalling matured memory B cells to sites shared between influenza virus subtypes (Ellebedy *et al.*, 2014; Nachbagauer *et al.*, 2014; Nachbagauer and Palese, 2020). Rational vaccine design of subunit-based vaccines is well-suited to this endeavor, as this approach uses structural insight of epitope:paratope interactions to produce a vaccine that induces specific and focused immune responses (Burton, 2010; Impagliazzo *et al.*, 2015; Yassine *et al.*, 2015; Kanekiyo *et al.*, 2019). Therefore, structural description of these conserved epitopes in complex with antibodies, as well as understanding the dynamics of the polyclonal antibody (pAb) response to these epitopes, is critical for rational universal influenza vaccine development.

Recent advances in evaluating antibody responses have produced a clearer picture of humoral immunity to influenza A virus (IAV), yet such techniques are still labor-intensive, time-consuming and unable to discern the full complexity of the pAb response (Wilson and Andrews, 2012). For example, enzyme-linked immunosorbent assays (ELISA) reveal antibody reactivity to immunogens but do not include information on targeted epitopes. Additionally, isolation and characterization of monoclonal antibodies (mAb) is a lengthy process involving multiple techniques and, due to limited sampling ability, typically only represents a subset of the pAb response. The application of single particle electron microscopy (EM) to structurally characterize heterogenous pAb immune complexes from low to high resolution yields unprecedented insight into polyclonal immune responses following IAV vaccination. We previously designed and implemented EM polyclonal epitope mapping (EMPEM) to discern the polyclonal immune response of rabbits and non-human primates to vaccination with HIV immunogens (Bianchi *et al.*, 2018; Cirelli *et al.*, 2019; Moyer *et al.*, 2020; Nogal *et al.*, 2020). This structure-based strategy is highly efficient: sample preparation is straightforward and the pipeline from sample collection to structural results is streamlined and expeditious, making longitudinal assessment of polyclonal responses in multiple subjects over a vaccination trial feasible.

Here, we employ EMPEM to map the landscape of human pAbs to HA after two doses of H5N1 vaccination (A/Indonesia/5/2005) from day 0 through 500, providing unique and comprehensive structural insights of polyclonal antibody responses against influenza virus. In a previous study that characterized H5N1-specific B cell responses of the same subjects in this trial, Ellebedy et al. determined that cross-reactive, stem-specific memory B cells respond to the first dose while head-specific, naïve B cells respond to the second dose of H5N1 vaccination (Ellebedy *et al.*, 2020, *in press*). Here, we elucidate the complexity of pAb responses to H5N1 vaccination through 500 days. We detect persistent pAb responses to heterosubtypic, broadly neutralizing epitopes on the stem domain of HA as well as pAb responses to the vulnerable sites on the head domain of HA that are also targeted during natural H5N1 infection in humans (Zuo *et al.*, 2015).

## RESULTS

### Robust serum response to H5N1 vaccination

To assess the humoral response to vaccination with a novel strain of influenza, we obtained serum from human subjects who participated in a pandemic H5N1 vaccination trial (NIH Clinical Trial Identifier: NCT01910519, A/Indonesia/5/2005 H5N1) (Ellebedy *et al.*, 2020, *in press*). The monovalent vaccine with Adjuvant System 03 (AS03), a squalene-in-water emulsion adjuvant, was administered in two immunizations at day 0 and day 21 (Figure 1A). Sera were collected at days 0, 7, 21, 28, 42 or 100, and 500 to measure the humoral response to H5N1 vaccination (Figure 1A).

**Figure 1.**
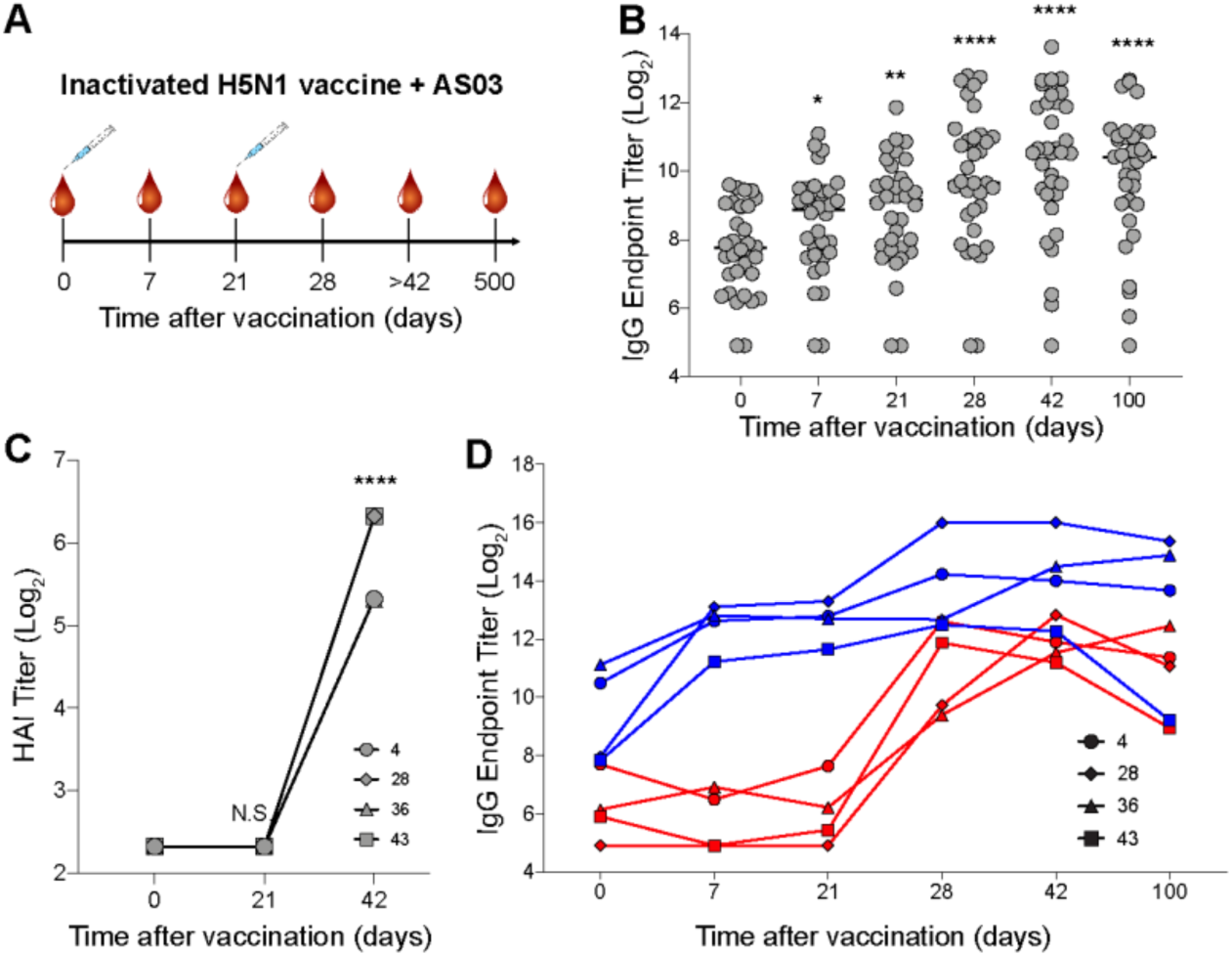
Novel H5N1 vaccination elicits robust IgG response to stem and head of hemagglutinin. (A) H5N1 vaccine trial: 34 healthy adults received the first immunization of inactivated H5N1 (A/Indonesia/5/2005) vaccine adjuvanted with AS03 at day 0 and the second immunization at day 21. Blood samples were collected on days 0, 7, 21, 28, 42 or 100, and 500 post vaccination. (B) ELISA binding titers of serum IgG to recombinant H5 HA over the course of vaccination. (C) HAI titers of serum IgG to recombinant H5 HA at days 0, 21, and 42 from subjects 4, 28, 36, and 43. (D) ELISA binding titers of serum IgG to the head or stem domains of H5 HA using H5 head-specific (red) and stem-specific (blue) probes from subjects 4, 28, 36, and 43. p-values were determined by unpaired t tests: * p-value <0.05, ** p-value < 0.01, *** p-value < 0.001, **** p-value < 0.0001, N.S. non-significant. See also Figure S1.

Previous work demonstrated that these subjects generated robust HA-specific plasmablast responses following H5N1 vaccination with AS03 adjuvant (Ellebedy *et al.*, 2020 *in press*). Through analysis of isolated mAbs we distinguished naïve and memory B cell dynamics of subjects 4, 28, 36, and 43 (Ellebedy *et al.*, 2020 *in press*). In the present study, we structurally characterized serum from these four subjects, with serological analyses to contextualize structural assessments. We observed a robust increase in H5-HA-specific serum IgG titers throughout the trial, with peak levels around 42 days after the first vaccination (Figure 1B and S1A). HAI titers, a measurement of the ability of HA to crosslink red blood cells through binding its receptor, sialic acid, increased after the second immunization in the four donors, suggesting that serum IgG at later time points neutralizes by blocking HA receptor binding activity (Figure 1C).

To dissect head- and stem-targeting pAbs in serum, we performed ELISA using probes of the trimeric HA head domain alone and a chimeric construct containing H5 stem and H9 head, as described previously (Ellebedy *et al.*, 2020 *in press*). Immediately following the first immunization, stem-specific serum IgG levels increased and remained high through 100 days post vaccination (Figure 1D). Conversely, head-specific serum IgG levels remained around baseline after the first immunization but rose drastically after the second immunization and remained elevated through 100 days post vaccination (Figure 1D). Monoclonal antibodies isolated from plasmablasts at days 7 and 28 from subjects 4, 28, 36, and 43 supported the head- and stem-specific dichotomy: day 7 mAbs targeted the stem while day 28 mAbs targeted the head of HA (Ellebedy *et al.*, 2020 *in press*). Affinity-matured B cells produce antibodies with high somatic hypermutation (SHM), long half-lives, and strong affinity, while naïve or new memory B cells produce antibodies with minimal mutations, short half-lives, and low affinity (Kurosaki, Kometani and Ise, 2015). We observed that day 7 mAbs had higher SHM loads (determined previously: Ellebedy *et al.*, 2020 *in press*). stronger affinities, and longer half-lives than day 28 mAbs (Figure S1B-E). These data suggest that affinity-matured memory B cells targeting the stem of HA responded quickly to novel H5N1 antigen but naïve or minimally-mutated memory B cells responded to the head of HA only after a second exposure to H5N1 antigen (Ellebedy *et al.*, 2020, *in press*).

While traditional serological analyses generate useful information about binding and functionality of pAb responses as a whole, they do not reveal details about epitopes and binding specificities of individual pAbs. Thus, we employed EMPEM to chart the epitope landscape targeted by pAbs over time.

### Polyclonal antibodies elicited by H5N1 vaccination bind the stem, head, and vestigial esterase domains of HA

EMPEM is a visual proteomics method that encompasses vaccination sample collection, antibody isolation and immune complex purification, and single particle EM image analysis (Figure 2). Following vaccination and antibody purification, we complex HA with a large molar excess of Fab and purify immune complexes by size exclusion chromatography, where a protein peak corresponding to the immune complex and unbound trimer separates from the excess Fab peak (Figure 2A and S2). During EM imaging and data processing, we collect micrographs, extract single particles, and categorize particles by similarity through multiple rounds of 2D classification (Figure 2B). In the 2D class averages, we can already detect numerous pAb specificities and assign targeted epitopes. Next, we perform multiple iterations of 3D classification and refinement of immune complexes. Finally, we represent 3D reconstructions of each pAb specificity on one protomer of the HA trimer to generate epitope landscapes of each subject and time point (Figure 2B).

**Figure 2.**
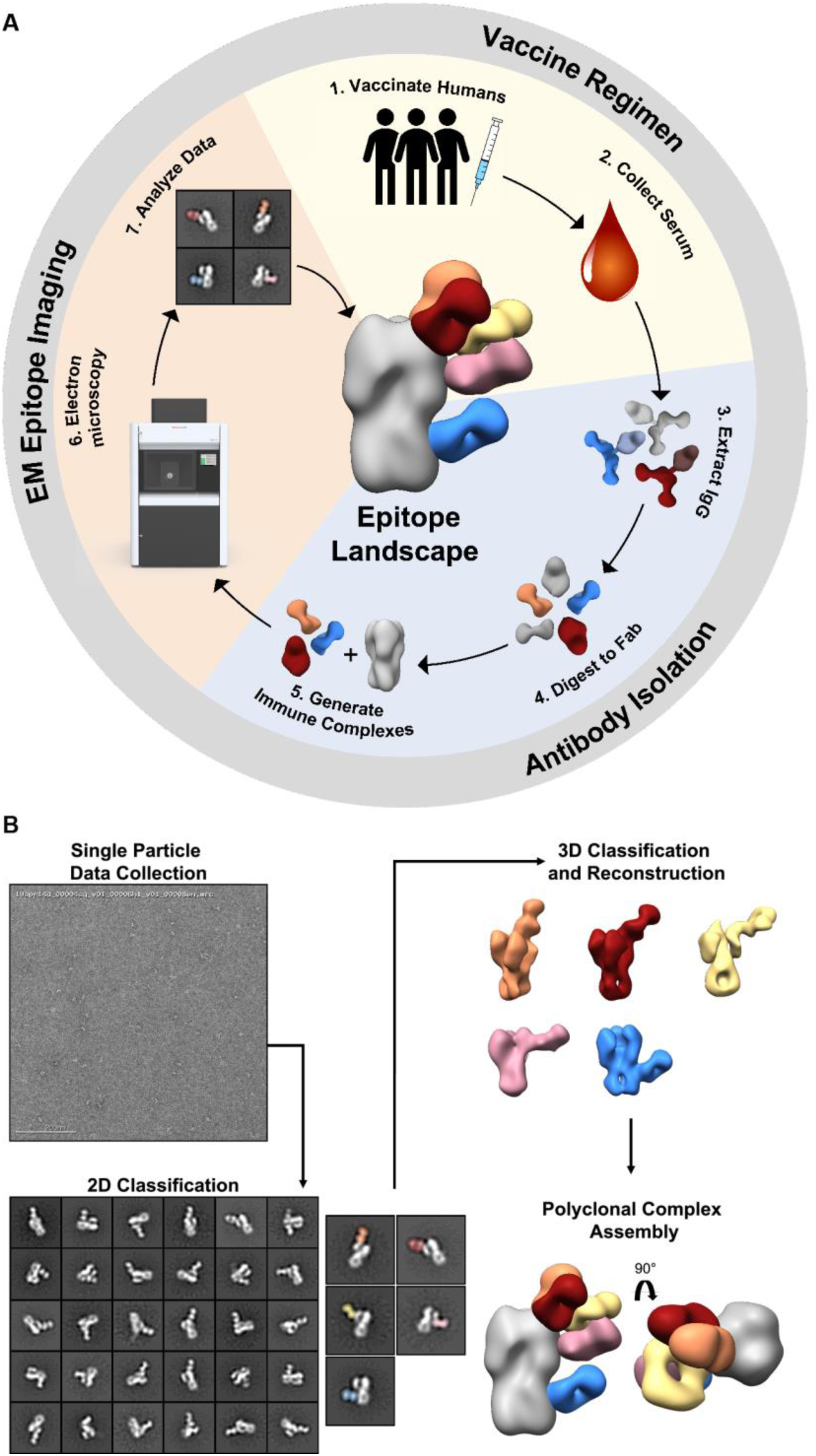
EM Polyclonal Epitope Mapping (EMPEM) generates epitope landscapes from single particles. (A) Overview of EMPEM technique. In the vaccine regimen stage human subjects are immunized (1) and serum samples collected throughout the course of the trial (2). During the antibody purification stage, IgG is extracted from serum samples (3) and digested with papain to Fab (4). Resulting Fabs are complexed with antigen (5). In the EM epitope imaging stage immune complexes are imaged by electron microscopy (6), single particle EM data are analyzed (7), and epitope landscapes are assembled. (B) Overview of EM data processing steps. Single particle EM data are collected and particles are categorized into 2D class averages. 2D classes of immune complexes are further categorized by another round of 2D classification and subjected to 3D classification and refinement. Finally, polyclonal complex assemblies are generated by segmenting and resampling densities corresponding to each Fab and mapped onto an HA trimer. See also Figure S2.

To characterize the dynamics of the pAb response to H5N1 vaccination, we performed EMPEM on serum samples from four subjects. For the first subject (subject 4), we processed sera from days 0, 7, 21, 28, 42, and 500 and complexed pAbs with the vaccine strain-matched HA (A/Indonesia/5/2005, Figures 3 and S3). At day 0, we did not observe any HA-specific pAbs in 2D class averages (Figure 3A). By day 7 and day 21, we observed stem-specific pAbs binding H5 HA (Figure 3B-C). As it was too early to detect pAbs from naïve B cells responding to H5N1 one week after vaccination, these stem-specific pAbs are most likely cross-reactive recalled antibodies elicited to conserved epitopes in the HA stem domain by prior seasonal influenza infection or vaccination. By day 28, one week after the second immunization, we observed pAbs still targeting the stem domain of HA as well as pAbs targeting the head domain at multiple angles and orientations around the receptor binding site (RBS), lateral patch, and vestigial esterase domain (Figure 3D). By day 42, pAb responses to the RBS, vestigial esterase domain, and stem domain persisted while other head-targeting pAbs waned (Figure 3E). Finally, after day 500, stem-specific pAbs remained circulating in low abundance (Figure 3F). Overall, subject 4 recalled cross-reactive stem-specific pAbs immediately following H5N1 vaccination that persisted over one year after vaccination and also elicited a transient wave of pAbs targeting the head and vestigial esterase domains after the second immunization.

**Figure 3.**
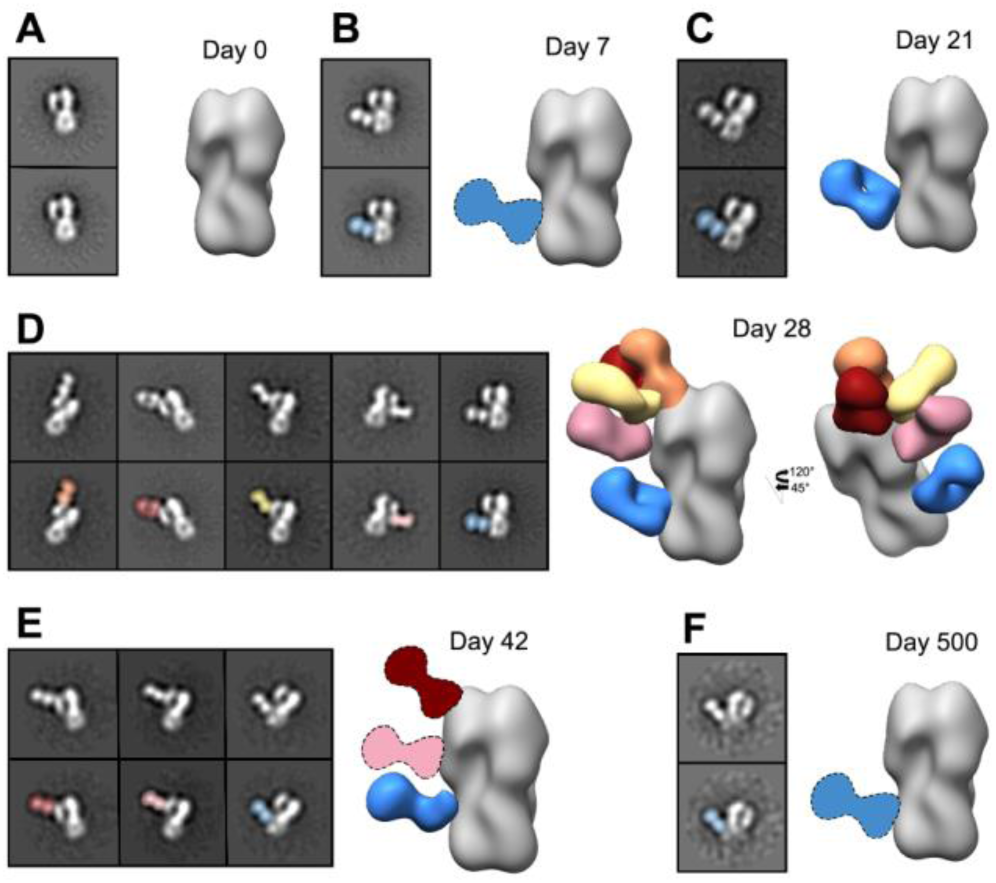
Kinetics of the polyclonal antibody response to H5N1 vaccination in subject 4. (A-F) Negative stain EM reconstructions of subject 4 polyclonal antibodies in complex with recombinant H5 HA (A/Indonesia/5/2005) at days 0 (A), 7 (B), 21 (C), 28 (D), 42 (E), and 500 (F) post vaccination. Top left panel: example 2D class average. Bottom left panel: 2D class average with the Fab labeled. Right panel: side view of polyclonal immune complexes. Stem specificity: blue, RBS-proximal and lateral patch specificities: red, orange, and yellow, vestigial esterase specificity: pink. Due to limited particle representation, Fab graphics with dashed outlines are predicted placements. See also Figure S3.

We extended EMPEM analyses to three more subjects in the H5N1 vaccination trial (Figures 4 and S4). At day 0, two of the subjects (36 and 43) already showed a stem-specific pAb response to H5 HA. These pAbs must therefore have been circulating in serum before H5N1 vaccination and cross-reacted with the vaccine antigen. By day 7, all subjects recalled a strong stem-specific memory pAb response to H5 HA (Figure 4A). These are likely broadly cross-reactive pAbs elicited by previous seasonal vaccination or infection. By day 28, all subjects maintained stem-specific pAb responses (Figure 4A) while eliciting head-specific pAb responses to varying degrees. Subjects 4 and 43 exhibited pAb responses to the RBS, lateral patch, and vestigial esterase domain. Subjects 28 and 36 had a less-diverse head-specific response to the RBS or to the mid-lateral region of the head. At intermediate time points (day 42 for subject 4 and day 100 for subject 43; serum unavailable for subjects 28 and 36 at these time points), pAbs targeting the RBS, vestigial esterase, and stem domains remained in circulation (Figure 4A). Expansion of pAbs against epitopes on the head of HA was consistent with the increase in HAI titers after the second immunization (Figures 4 and 1C). After 500 days all subjects with serum available demonstrated a lasting stem-specific pAb response (Figure 4), and subject 28 also displayed an RBS-specific pAb response.

**Figure 4.**
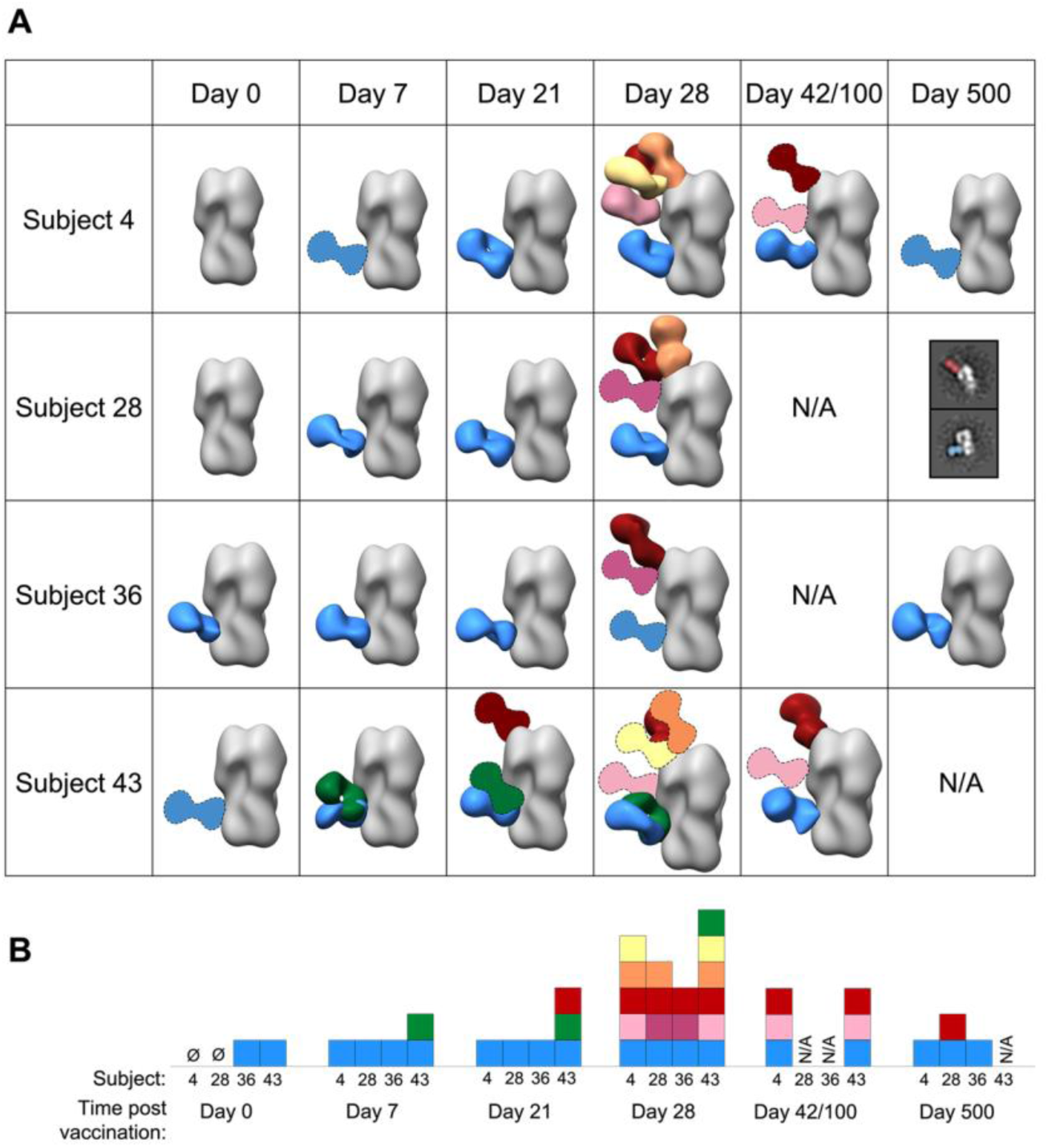
Polyclonal antibodies elicited by H5N1 vaccination decorate the stem, head, and vestigial esterase domains of HA. (A) Matrix of negative stain EM reconstructions of polyclonal antibodies in complex with recombinant H5 HA (A/Indonesia/5/2005) from each subject at all time points listed. 3D reconstructions of polyclonal immune complexes are shown for the majority of time points. Due to limited particle representation, Fab graphics with dashed outlines are predicted placements. For samples with immune complexes in low abundance, example 2D class averages with labels are shown. (B) Summary of epitopes targeted by polyclonal antibodies. Each square represents a Fab specificity from the corresponding subject and time point. Stem specificities: blue and green, RBS-proximal and lateral patch specificities: red, orange, and yellow, vestigial esterase and mid-lateral head specificities: light pink and dark pink. See also Figure S4.

In summary, the landscape of pAbs responding to H5N1 vaccination followed two concurrent trends: stem-specific pAbs targeted H5 HA from baseline through 500 days (Figure 4B). These pAbs may have been circulating in serum before vaccination, recalled shortly after the first immunization, or newly elicited after the second immunization. Head-specific pAbs expanded after the second immunization, targeting the RBS, lateral patch, mid-lateral head region, and vestigial esterase region, and waned quickly as the trial progressed, with the pAbs targeting more conserved regions enduring longer (Figure 4B). Together, these data demonstrate that H5N1 vaccination elicits a prominent and prolonged pAb response to the conserved stem domain of HA, as well as a more diverse and transient pAb response to the variable, immunodominant head domain of HA.

The kinetics of the polyclonal response mapped by EM are consistent with serum ELISA and HAI data showing that stem-specific antibodies were already present at day 0 and continually increased over the course of vaccination, while head-specific antibodies and HAI+ antibodies were substantially present only after the second immunization (Figure 4 and 1C-D). By day 42 or 100, serum IgG binding titers were still high for the majority of subjects but were declining for donors 4 and 43 (Figure 1B and D). In agreement, the EM-mapped pAb landscape at these intermediate time points was less diverse for donors 4 and 43. Perhaps more dominant pAbs persist at high levels through these time points while the more transient pAbs wane.

### H5N1 vaccine-elicited mAbs target discrete regions on HA and share footprints with pAbs

Previously, Ellebedy et al characterized a panel of mAbs isolated from these subjects and identified a biphasic response: day 7 mAbs exclusively targeted the stem of HA with high SHM and breadth while day 28 mAbs targeted the head of HA with little SHM and strain or subtype - specific reactivity (Ellebedy *et al.*, 2020 *in press*). Here, we complexed mAbs at days 7 and 28 from subjects 4 and 43 with H5 HA (A/Indonesia/5/2005) to compare the binding footprints and orientations with pAbs at day 28. All day 7 mAbs used the V_H_ 1-69 germline and targeted the same footprint on the stem of HA (Figures 5A and S5A-B). While day 28 mAbs utilized a mixture of germlines, including three mAbs with V_H_ 3-33, six of the seven mAbs targeted the same footprint on the RBS of HA (Figures 5C, S5A, and S5C). The seventh mAb targeted the vestigial esterase domain of HA (Figure 5B). Upon comparing the footprints of these mAbs with day 28 pAbs from subjects 4 and 43, we observed that each day 7 mAb overlapped almost perfectly with the blue stem-specific pAbs (Figure 5D). Additionally, each day 28 RBS-specific mAb overlapped almost exclusively with each other and were represented by pAbs (Figure 5D). Finally, while we were unable to obtain a 3D reconstruction of day 28 1B02 mAb, 2D class averages confirmed that it bound the vestigial esterase domain in the same vicinity as the vestigial esterase-specific pAb (Figures 5B and 5D).

**Figure 5.**
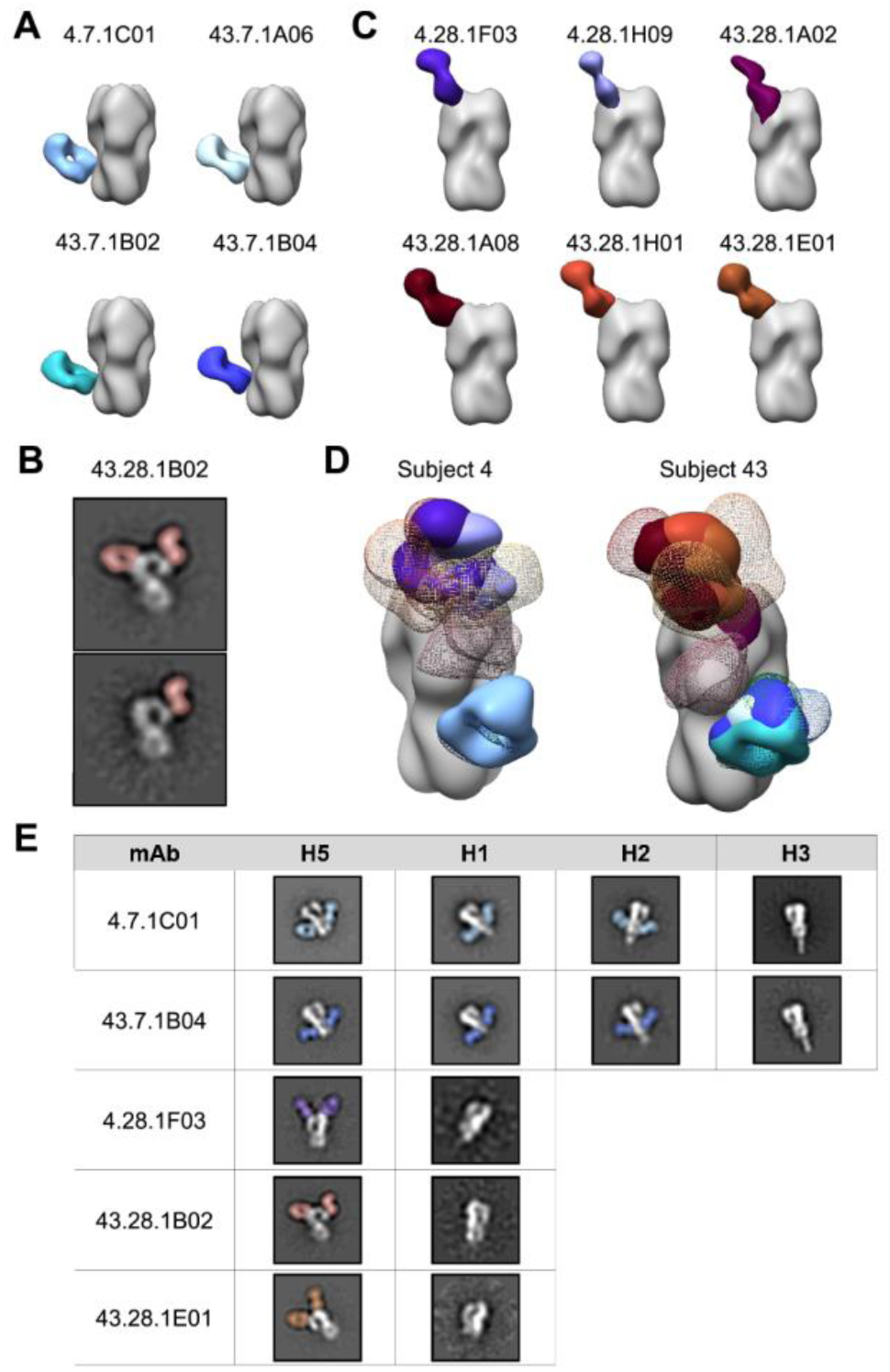
H5N1 vaccine-elicited mAbs target discrete regions on HA and share footprints with polyclonal antibodies. (A-C) Side views of negative stain EM reconstructions or 2D class averages show subjects 4 and 43 day 7 mAbs target the stem domain (A) while day 28 mAbs target the vestigial esterase domain (B) and the RBS (C) of recombinant H5 HA (A/Indonesia/5/2005). (D) Comparison of subjects 4 and 43 mAb and pAb immune complexes. Due to limited particle representation, orange, yellow, and pink pAbs from subject 43 are predicted placements. (E) Cross-reactivity of example mAbs from subjects 4 and 43 to recombinant HAs from A/Indonesia/5/2005 (H5N1), A/California/04/2009 (H1N1), A/Singapore/1/1957 (H2N2), and A/Singapore/INFIMH-16-0019/2016 (H3N2). mAbs are shown in solid colors; 4.7.1C01 pale blue, 43.7.1A06 light blue, 43.7.1B02 teal, 43.7.1B04 dark blue, 4.28.1F03 purple, 4.28.1H09 lavender, 43.28.1A02 plum, 43.28.1A08 maroon, 43.28.1H01 brick, and 43.28.1E01 brown. pAbs from day 28 are shown in colored mesh; stem specificities: blue and green, RBS-proximal and lateral patch specificities: red, orange, and yellow, vestigial esterase and mid-lateral head specificities: pink. See also Figure S5.

To compare the cross-reactive potential of mAbs from days 7 and 28, we complexed each mAb with HAs from heterosubtypic strains including H1N1 (A/California/04/2009), H2N2 (A/Singapore/1/1957), and H3N2 (A/Singapore/INFIMH-16-0019/2016; Figure 5E). As expected, day 7 stem-specific mAbs demonstrated broad cross-reactivity, binding H5, H1, and H2 HAs, while day 28 head-specific mAbs demonstrated subtype- or strain-specific reactivity, binding only H5 HA.

Out of six pAb footprints identified by EMPEM only three were represented by mAbs, suggesting that mAb analysis did not fully recapitulate the diversity of the polyclonal response. Additionally, while day 28 mAbs targeted the head of HA almost exclusively (Figure 5C-D), ELISA data of serum IgG demonstrate equal or stronger binding activity to the stem than the head of HA at day 28 (Figure 1D), further highlighting the role of pAb mapping to elucidate the complete polyclonal response at the serum level. However, mAb data provide valuable functional information and are complementary with EMPEM data; taken together these results suggest that pAbs at day 7 are dominated by robust and broadly cross-reactive V_H_ 1-69 stem responses while pAbs at day 28 target variable head epitopes and continue to target broadly cross-reactive stem epitopes.

### Heterosubtypic immunity by H5N1 vaccine-elicited polyclonal antibodies

After characterizing polyclonal epitope landscapes at low resolution, we next aimed to decipher molecular details of epitope:paratope interactions through cryoEMPEM on polyclonal sera. As serum samples were already depleted from our previous study as well as negative stain EMPEM, we only had enough serum to perform this analysis on pAbs from subject 4 at day 28. We processed cryoEM data using a focused classification data analysis pipeline (Figure S6A) (Zivanov *et al.*, 2018) that enabled detection and reconstruction of minority immune complexes from total particles.

We discerned high resolution immune complexes of pAbs from subject 4 at day 28 that targeted the stem and multiple sites on the head of HA from a single cryoEM sample (Figures 6A-B and S6B-E). Importantly, these epitopes corroborated the negative stain epitope landscape for subject 4 at day 28, demonstrating that the lower resolution negative stain methodology was sufficient to observe all epitopes. Due to limited serum sample availability each immune complex in our cryoEM images was present in low abundance—the stem specificity accounted for ∼4% while each of the three head specificities represented just ∼1-2% of the total particles. Nevertheless, focused classification enabled us to detect and reconstruct these antibody:epitope interactions. The RBS-binding pAb plugged the receptor binding pocket on HA, likely blocking receptor binding (red epitope, Figure 6A-B and Figure S6B). The remaining head-binding pAbs targeted the lateral patch (yellow) and vestigial esterase (pink) epitopes (Figure 6A-B and Figure S6B).

**Figure 6.**
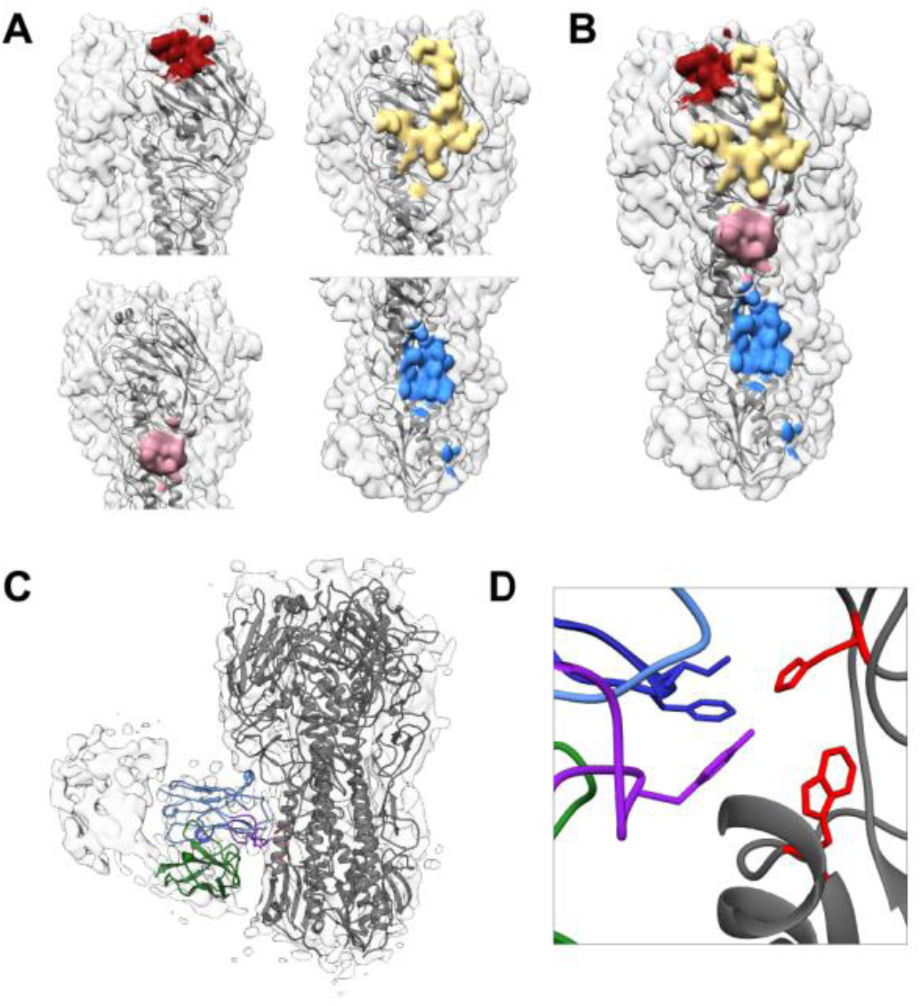
High resolution cryoEM defines broadly reactive epitopes of H5N1 vaccine-elicited polyclonal antibodies. (A) CryoEM-mapped epitopes targeted by pAbs from subject 4 at day 28. Red: RBS epitope. Yellow: lateral patch epitope. Pink: vestigial esterase epitope. Blue: V_H_1-69 epitope. (B) Each epitope specificity marked together on a single trimer. (C-D) CryoEM map of stem immune complex from subject 4 at day 28 with ribbon diagrams for H5 HA (grey; A/Indonesia/5/2005; PDB 4K62) and mAb 1C01 from subject 4 day 7 docked into the EM density. Full length side view (C) and zoomed in view of the epitope:paratope interaction (D). V_H_: blue, V_L_: green, CDRH2: dark blue, CDRH3: purple, HA residues H18 (top) and W21 (bottom): red. See also Figure S6.

Intriguingly, the stem-specific pAb (Figure 6A-C) bound in almost the exact same location and orientation as CR9114, the gold standard stem-directed broadly neutralizing Ab (Figure S6F) (Dreyfus *et al.*, 2012; Lee and Wilson, 2015). The V_H_ 1-69 lineage broadly reactive mAb 1C01 from subject 4 superimposes almost perfectly with the stem-binding pAb, suggesting that this mAb is part of the polyclonal response at that specificity (Figure 5A and S6G). Upon docking the predicted structure of the variable regions of 1C01 mAb into the cryoEM pAb density, we observed that the pAb interaction appears to be mediated by the signature IFY motif in CDR H2 and H3 that likely interacts with H18 in HA1 and W21 in HA2 (Figure 6C-D) (Dreyfus *et al.*, 2012; Lee and Wilson, 2015). Thus, high resolution cryoEM mapping identified a broadly neutralizing CR9114-like stem response in polyclonal sera following H5N1 vaccination, supporting the novel H5 antigen as a prime candidate for a universal influenza vaccine.

The major vulnerable sites (VS) on the head of H5 HA have been determined previously based on protective neutralizing mAbs isolated from humans who recovered from HPAI H5N1 infections (VS1-3) and mice immunized with HPAI H5N1 (VS4) (Zhu *et al.*, 2013; Zuo *et al.*, 2015). We mapped these vulnerable sites onto the H5 HA trimer and compared them to negative stain EMPEM epitopes (Figure 7A). The footprints of H5N1 vaccine-elicited pAbs determined by both negative stain and cryoEM encompassed each of the vulnerable sites, including the RBS in VS2 (red), lateral patch in VS1 (yellow), and vestigial esterase domain bridging VS3 and 4 (pinks; Figures 6A-B and 7A). The major vulnerable sites were determined based on protective neutralization activity. Though antibody responses to the stem can have neutralizing activity, they can also protect through effector functions such as antibody-dependent cell-mediated cytotoxicity and antibody-dependent cellular phagocytosis (Dilillo *et al.*, 2014; Cox *et al.*, 2016; Coughlan and Palese, 2018; Boudreau and Alter, 2019). H5N1 vaccine-elicited pAbs also targeted the stem domain, but were not represented in the characterization of neutralizing responses to H5N1 infection (Zuo *et al.*, 2015). In summary, H5N1 vaccine-elicited head pAb responses mimic broadly neutralizing mAb responses to natural HPAI H5N1 infection.

**Figure 7.**
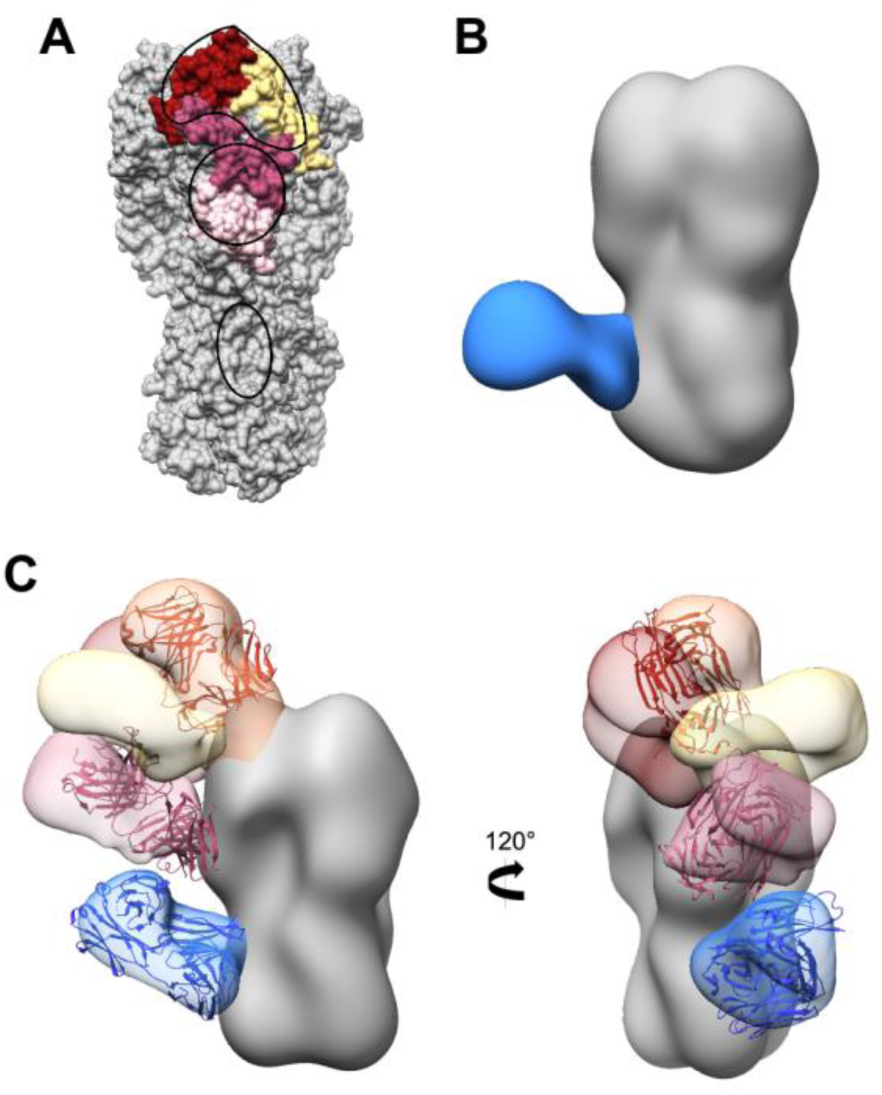
H5N1 vaccine-elicited pAb response mimics natural infection and generates heterosubtypic immunity. (A) H5 HA (A/Indonesia/5/2005) with vulnerable sites (VS) marked in yellow for VS1, red for VS2, dark pink for VS3, and light pink for VS4. Footprints of pAbs from all subjects at day 28 are outlined in black. (B) Negative stain EM reconstruction of subject 43 polyclonal antibodies in complex with recombinant H1 HA (A/Michigan/45/2015) at day 21. Stem specificity: blue. (C) Subject 4 day 28 negative stain epitope landscape with broadly neutralizing antibodies docked into HA density. Orange: bnAb S139/1 (PDB 4GMS), pink: bnAb H5M9 (4MHH), blue: bnAb CR9114 (4FQI). See also Figure S7.

To investigate the heterosubtypic breadth of polyclonal antibodies elicited by H5N1 vaccination, we examined the ability of pAbs from donor 43 at day 21 to bind H1 HA using EMPEM (Figure 7B and S7). We assessed cross-reactivity to the antigenically drifted A/Michigan/45/2015 (H1N1) HA that emerged in 2015. As serum samples were isolated from subjects in 2013, the subjects were naïve to this H1N1 strain. Due to limited serum sample availability, we were only able to perform this analysis on one time point. However, we saw pAbs targeting the broadly cross-reactive V_H_ 1-69 epitope on the stem of H1 HA, suggesting that H5N1 vaccination elicits pAbs that recognize multiple influenza virus subtypes. Moreover, these results demonstrate that EMPEM can sensitively detect pAb responses to strains of different subtypes than the vaccine strain.

To compare the epitope landscapes after H5N1 vaccination with known broadly neutralizing responses, we superimposed H5N1-reactive broadly neutralizing antibodies (bnAbs) with pAbs from subject 4 at day 28 (Figure 7C) (Sun *et al.*, 2014; Lee and Wilson, 2015). As described above, the bnAb CR9114 that protects against both influenza A and B viruses mapped almost exactly with vaccine-elicited stem pAbs and V_H_ 1-69 mAbs (Figures S6F-G and 7C) (Dreyfus *et al.*, 2012). Similarly, bnAbs toward the vestigial esterase (H5M9, pan-H5N1 reactive) and RBS (S139/1, group 1 and 2 cross-reactivity against H1, H2, H3, H5, H9, and H13) also overlapped footprints and angles of approach with pAbs (Figure 7C) (Yoshida *et al.*, 2009; Lee *et al.*, 2012; Zhu *et al.*, 2013). These results suggest that H5N1 vaccination elicits pAb responses to regions of broadly neutralizing epitopes.

## DISCUSSION

Comprehensive mapping of the HA epitopes targeted by the polyclonal antibody response remains a major gap in our understanding of the human B cell response to influenza viruses. EMPEM is a unique modality that sensitively detects minority antigen-specific antibodies, providing a comprehensive landscape of the polyclonal immune response. In this study, we show that EMPEM can directly inform rational design of a universal influenza vaccine by extensive characterization of pAb responses to a vaccine derived from non-circulating IAV strains, such as H5N1. While we have performed EMPEM on rabbit and NHP sera, here we report the first study using our structure-based method to comprehensively map the polyclonal response to an avian influenza virus vaccine in humans (Bianchi *et al.*, 2018; Cirelli *et al.*, 2019; Moyer *et al.*, 2020; Nogal *et al.*, 2020).

Complementary serological techniques inform our understanding of the pAb response to influenza virus vaccination, and EMPEM contextualizes data from these traditional methods. Here, ELISA binding titers indicated a strong stem-specific response that increased over the course of H5N1 vaccination but a low initial head-specific response that expanded after the second immunization (Figure 1). HAI+ antibodies were only detected after the second immunization (Figure 1). Ellebedy et al. also determined that mAbs at day 7 had higher levels of somatic hypermutation and were likely generated by highly mutated memory B cells, while mAbs at day 28 were strain-specific and were produced from naïve or recently generated memory B cells (Ellebedy *et al.*, 2020 *in press*).

EMPEM analyses corroborated and extended these observations by describing specific epitopes targeted on the head and stem domains of HA, differentiating overlapping antibodies and angles of approach to similar epitopes, and monitoring the kinetics of each response over the course of vaccination (Figures 3-4). EMPEM analyses also elucidated more intricate details including persistent, heterosubtypic stem pAbs with V_H_ 1-69 CR9114-like qualities and head pAbs that mirrored responses to natural H5N1 infection (Figures 6-7). Additionally, the number of unique specificities in the pAb response began declining by day 42 or 100 in two donors, supported by ELISA data also showing declining serum IgG titers for these donors (Figure 4 and 1D). These results demonstrate that EMPEM can track the kinetics of individual pAb specificities. Interestingly, subject 28 still had low levels of pAbs targeting the RBS region circulating one year after vaccination (Figure 4). As these pAbs may recognize conserved aspects of the RBS, they may have been derived from a new memory B cell population that matured and differentiated into long-lived plasma cells. In the future, we can test this hypothesis by tracking the evolution of RBS-specific mAbs from subject 28 over the course of the trial.

By matching binding specificities of mAbs and pAbs from corresponding serum samples, we can evaluate how well mAbs represent the pAb response and extrapolate functional properties of pAbs. Monoclonal Abs are usually isolated from plasmablasts or memory B cells and thus may not encompass the diversity of antibodies present in serum. Here, mAb analyses demonstrated a biphasic response to H5N1 vaccination; mAbs isolated at day 7 exclusively bound the stem domain while mAbs from day 28 specifically bound the head domain of HA. Interestingly, the biphasic response to HA was not fully recapitulated in EMPEM and ELISA data; a large proportion of pAbs still targeted the stem after the second immunization (Figures 1D, 3, and 4). Moreover, EM mapped day 28 mAbs bound almost exclusively to the RBS, while day 28 EMPEM results demonstrated a diverse pAb response to multiple head epitopes (Figures 4-5) (Ellebedy *et al.*, 2020 *in press*). We used full length H5 HA as the probe for isolating mAbs; perhaps BCRs recognize the RBS more readily than the stem resulting in biased isolation of head-specific B cells. Alternatively, even though the quantity of stem pAbs did not diminish, it is possible that the expansion of head-specific B cells at day 28 outnumbered stem-specific B cells, resulting in preferential isolation of head-specific B cells. Finally, due to clonal expansion, mAb isolation may be biased toward more dominant B cell clones, whereas EMPEM evaluates the total IgG pool and can identify minority epitopes. Nevertheless, we can extract crucial details about the polyclonal response through mAb analysis; day 7 pAbs consist of broadly reactive V_H_ 1-69 antibodies. Monoclonal Ab analysis provides powerful functional details of antibodies but EMPEM is necessary to recapitulate the complexity and dynamics of the pAb response.

Humans have lifelong exposure history with influenza viruses and elicit immune responses mostly to the immunodominant and drifting head of HA (Krammer, 2019). An individual’s first influenza virus infection strongly biases their immune response to subsequent, drifted influenza virus strains, resulting in person-to-person variation (Li *et al.*, 2013; Andrews *et al.*, 2015; Gostic *et al.*, 2016, 2019). Additionally, single Ab specificities may functionally dominate the pAb response, rendering other specificities inconsequential (Li *et al.*, 2013; Huang *et al.*, 2015; Lee *et al.*, 2019). This phenomenon also differs between individuals and may be influenced by pre-exposure history. Our study establishes EMPEM as a powerful new tool for gauging pre-existing immunity and variation between individuals by mapping bulk influenza-directed pAb responses at baseline and over the course of vaccination to multiple antigens as well as distinguishing differences in epitope targets per individual. Our EMPEM results identified pre-existing, cross-reactive stem pAbs at baseline in half of the subjects, elicitation and persistence of cross-reactive stem pAbs in all subjects after primary exposure, and expansion of a decorated head response that varied in each subject after secondary exposure to H5 HA (Figures 3-4). In future studies with known patient ages and immune histories, EMPEM will be advantageous for dissecting pAb targets between individuals, connecting pAb differences with pre-exposure immune profiles.

As humans are immunologically naïve to avian influenza virus strains, such as H5N1, primary exposure to their antigens may direct the immune response to conserved epitopes predominately located on the stem region of HA, potentially providing broad protection against circulating and pandemic influenza virus strains (Nachbagauer and Palese, 2020). Previous studies suggested that H5N1 vaccines are poorly immunogenic and result in lower seroconversion rates compared to seasonal influenza virus vaccination, but these conclusions are mostly based off of HAI titers (Bresson *et al.*, 2006; Treanor *et al.*, 2006; Belshe *et al.*, 2014). Here we show that low dose inactivated H5N1 vaccination adjuvanted with AS03 elicits a potent response to highly conserved epitopes in the stem domain immediately after the first immunization that sustains over a year after vaccination. Additionally, after the second immunization we observed a head-specific response that targets all major neutralizing sites on the head of H5 HA, potentially enhancing protection against H5N1 strains (Figure 7A) (Zuo *et al.*, 2015). Finally, H5N1-elicited pAbs encompass broadly neutralizing sites in the stem, esterase, and RBS of HA, emphasizing H5 HA’s utility as an antigenic candidate (Figure 7C).

Training adaptive immunity to successfully target and neutralize multiple influenza virus strains is imperative for the success of a universal influenza vaccine. This study underscores the ability of novel influenza antigens to elicit memory responses to conserved sites on HA, while also illustrating subtype-specific responses upon re-exposure to the same strain. Therefore, immunogens in a prime/boost regimen for a universal influenza vaccine should be from a different strain, such as a novel H7 antigen, to keep pAb responses trained on conserved epitopes and away from off-target recalled epitopes. Here and in future studies, EMPEM can dissect pAb responses to influenza virus vaccinations and infections with novel and seasonal strains, enabling comparison of immune responses of humans across age groups and with varying exposure histories. This extensive characterization of immune responses to influenza virus will contribute invaluable information for developing universal influenza vaccines.

## ACKNOWLEDGEMENTS

Recombinant HA from A/Indonesia/5/2005 (H5N1) was provided by the International Reagent Resource. We thank Aleksandar Antanasijevic for invaluable assistance with focus classification data processing, Ian Wilson, Adrian McDermott, and James Crowe for providing recombinant HA protein, Bill Anderson for keeping the EM suite at Scripps Research running efficiently, and Alba Torrents de la Peña and Lauren Holden for critical reading of the manuscript. The Ward laboratory was funded by Collaborative Influenza Vaccine Innovation Centers contract 75N93019C00051. Julianna Han was funded by NIAID 2 T32 AI007244-36. The Ellebedy laboratory was supported by NIAID grants R21 AI139813, U01 AI141990, and NIAID Centers of Excellence for Influenza Research and Surveillance (CEIRS) contract HHSN272201400006C.

## AUTHOR CONTRIBUTIONS

JH, ABW, AHE, and RA conceptualized the project. JH designed and performed EMPEM experiments, microscopy, and data processing. AHE performed serological studies and advised on the project. AJS produced mAbs. YND performed BLI and SPR experiments and analyzed data. STR, HLT, and BMM assisted with EMPEM experiments and microscopy. DF and RA advised on the project. ABW supervised the project. JH wrote the manuscript, ABW edited the manuscript, and all authors provided feedback for the manuscript.

## METHODS

### RESOURCE AVAILABILITY

#### Lead Contact

Further information and requests for resources and reagents should be directed to and will be fulfilled by the Lead Contact, Andrew B. Ward.

#### Materials Availability

This study did not generate new unique reagents.

#### Data and Code Availability

3D EM reconstructions have been deposited to The Electron Microscopy Data Bank (emdataresource.org) under accession numbers listed in the Key Resources Table.

### EXPERIMENTAL MODEL AND SUBJECT DETAILS

The vaccination trial (NIH Clinical Trial Identifier: NCT01910519) has been described previously and was approved by the Institutional Review Board of Emory University (Ellebedy *et al.*, 2020 *in press*). Briefly, healthy adult male and female participants between 21-45 years old were immunized with monovalent inactivated A/Indonesia/05/2005 (H5N1) influenza vaccine with AS03 adjuvant provided by GlaxoSmithKline. All participants provided informed consent. Plasma and peripheral blood mononuclear cells (PBMCs) were collected at days 0, 7, 21, 28, 42, 100, and 500 after the first immunization.

## METHOD DETAILS

### ELISA

We coated 96-well plates with recombinant HA (A/Indonesia/5/2005) overnight at 4°C. Following HA binding, we incubated the plates at room temperature for 1 hour with blocking buffer (0.1% Tween 20, 0.5% milk powder, 3% goat serum in phosphate-buffered saline (PBS)). Next, we incubated serial dilutions of mAbs on HA-coated plates at room temperature for 2 hours followed by three washes with 0.1% Tween 20 in PBS. We added secondary goat anti-human IgG conjugated to horse radish peroxidase at 1:3000 dilution in blocking buffer to plates and incubated at room temperature for 1 hour. Finally, we washed the plates four times with 0.1% Tween 20 in PBS, added SigmaFast o-phenylenediamine solution, and measured 490nm signal using a plate reader. To distinguish head and stem responses, we used the trimeric head domain from A/Indonesia/5/2005 and chimeric HA expressing the stem from A/Indonesia/5/2005 and the head of H9 from A/guinea fowl/Hong Kong/WF10/1999 (H9N2), produced recombinantly using the baculovirus expression system described previously (Ellebedy *et al.*, 2014).

### HA inhibition

We added serially diluted mAbs at an initial concentration of 30μg/mL in PBS to V-shaped 96-well plates in duplicate. Next, we added 8 HA units/50μL of 6:2 re-assortant, low pathogenic (no multi-basic cleavage site) H5N1 virus (A/Indonesia/05/2005 and PR8 IBCDC-RG (H5N1)) to each well and incubated plates at room temperature for 30 min with shaking. We added 0.5% chicken red blood cells to each well and incubated plates at 4°C until red blood cells formed puncta at the bottom of the negative control wells. We measured minimum HAI concentration by determining the last dilution well that did not display hemagglutination.

### mAb production

Antibodies were cloned as previously described (Wrammert *et al.*, 2008). Total or H5 HA probe-binding plasmablasts were single cell sorted into 96-well plates containing RNA stabilizing buffer. VH, Vκ, and Vλ genes were then amplified by reverse transcription-PCR and nested PCR reactions from singly-sorted GC B cells and PBs using cocktails of primers specific for IgG, IgM/A, Igκ, and Igλ from previously detailed primer sets and then sequenced. The amplified VH, Vκ, and Vλ genes were cloned into IgG1 and Igκ expression vectors, respectively, as previously described. Heavy and light chain plasmids were co-transfected into Expi293F cells (Gibco) for expression and antibody was purified with protein A agarose (Invitrogen).

### Affinity measurements

To measure binding affinity of HA to Fabs, we first digested mAbs to Fab with immobilized papain (Thermo Fisher) and purified the digestion products over a protein A/G affinity column to isolate Fab. We biotinylated HA in vitro (EZ-Link-NHS-PEG4-Biotin, Thermo Fisher) and removed excess biotin with a desalting column (0.5mL Zeba Spin 7K MWCO, Thermo Fisher). We measured binding affinity of each Fab to HA by bio-layer interferometry (BLI) using an Octet Red96 instrument (ForteBio) or surface plasmon resonance (SPR) using a Biacore T200 SPR instrument (GE Healthcare). Using BLI, we performed a preliminary experiment to identify Fabs with too low signal-to-noise ratio to quantify; for higher sensitivity, we conducted SPR analyses for these Fabs. For BLI, we loaded 5μg/mL biotinylated HA in HBS-EP buffer (10 mM HEPES pH 7.4, 150 mM NaCl, 3 mM EDTA, and 0.005% P20 surfactant) with 1% BSA onto streptavidin biosensors (ForteBio) for 1min. For SPR, we coupled neutravidin protein to CM5 sensor chips using standard amine-coupling in HBS-EP buffer and captured biotinylated HA, which resulted in a response of 200-300 RU. We measured Fabs in 3-fold serial dilutions at 25°C and processed data using Biaevaluation 3.1 (GE Healthcare). We employed a 1:1 binding model to measure association and dissociation rate constants and fit steady-state equilibrium concentration curves. We calculated t_1/2_ for each Fab using the formula t_1/2_=ln2/k_d_.

### Serum IgG purification, digestion, and complexing

To purify IgG from serum samples we heat inactivated 1 mL serum samples in a 55°C water bath for 30 min and incubated them on protein G resin (GE Healthcare) or Capture Select in a 1:1 ratio of serum to resin for 20 hours to bind IgG. After incubation, we removed IgG-depleted serum and washed IgG-bound protein G samples three times with PBS using centrifugation and Amicon concentrators. Next, to elute IgG from protein G resin, we incubated the samples in 0.1M glycine pH 2.5 buffer for 20 min followed by neutralization with 1M Tris-HCl pH 8 buffer, repeated twice. We buffer exchanged the samples into PBS using centrifugation with Amicon concentrators. For IgG digestion, we incubated 4 mg IgG with immobilized papain (Thermo Fisher) in freshly-prepared digestion buffer (20mM sodium phosphate, 10 mM EDTA, 20 mM cysteine, pH 7.4) at 37°C for 18-22 hours. We separated digested IgG and immobilized papain using Pierce spin columns (Thermo Fisher) and buffer exchanged digested IgG into tris-buffered saline (TBS) using centrifugation with Amicon concentrators. To separate Fab/Fc from IgG we ran the samples through size exclusion chromatography with a Superdex 200 increase 10/300 column (GE Healthcare). For complexing, we concentrated 1mg Fab/Fc to ∼50μL and incubated with 20μg recombinant HA at room temperature for 16-20 hours. Finally, we purified immune complexes from unbound Fab/Fc through size exclusion chromatography with a Superose 6 increase 10/300 column (GE Healthcare) and concentrated the immune complexes to ∼50μL.

### mAb digestion and complexing for EM

To digest monoclonal IgG to Fab, we first incubated papain in freshly-prepared digestion buffer (100mM Tris pH 8, 2mM EDTA, 10mM L-cysteine) at 37°C for 15min. Next, we incubated activated papain with 1mg IgG in freshly-prepared digestion buffer at 37°C for 2 hours. To end the digestion, we added 50mM iodoacetamide. We buffer exchanged the digestion products into PBS using centrifugation with Amicon concentrators and purified Fabs using a CaptureSelect CH1-XL column (Thermo Fisher), eluting Fab with 150 mM sodium chloride and 20mM sodium acetate pH 3.4 buffer and neutralizing with 1M Tris-HCl pH 8 buffer. We buffer exchanged and concentrated Fab into PBS using centrifugation with Amicon concentrators and complexed Fab with HA at greater than 3x molar ratio of Fab to HA at 4°C for 16-20 hours.

### Negative stain electron microscopy

To prepare grids for negative stain electron microscopy, we deposited purified immune complexes at ∼30μg/mL onto glow-discharged, carbon-coated 400 mesh copper grids (Electron Microscopy Sciences, EMS). After blotting to remove excess sample, we stained the grids with 2% w/v uranyl formate for 30s followed by blotting to remove excess stain. We imaged the grids on a Talos 200C with a Falcon II direct electron detector and a CETA 4k camera (FEI) at 200kV and 73,000x magnification, a Tecnai Spirit T12 (FEI) with a CMOS 4k camera (TVIPS) at 120kV and 52,000x magnification, and a Tecnai T20 (FEI) with an Eagle CCD 4k camera (FEI) at 200kV and 62,000x magnification. We collected micrographs using Leginon, picked and stacked 100,000-400,000 single particles using Appion, and processed particles to reference-free 2D class averages and 3D reconstructions using Relion (Suloway *et al.*, 2005; Lander *et al.*, 2009; Scheres, 2012; Zivanov *et al.*, 2018). We used UCSF Chimera to analyze data and generate figures (Pettersen *et al.*, 2004). Due to low angular sampling induced by particle orientation bias and low abundance of immune complexes, a small proportion of immune complexes only partially reconstructed in 3D but clearly showed Fab placement relative to HA. These placements were also confirmed by distinct 2D class averages. Similarly to Gilchuk et al., we used our internal database of HA immune complexes, partial 3D density, and distinguishing 2D class averages as references to project these specificities onto 3D models of HA and mark them as predicted placements (Gilchuk *et al.*, 2019).

### CryoEM

To freeze grids for cryoEM, we added Lauryl Maltose Neopentyl Glycol (Anatrace) at a final concentration of 5μM to purified immune complexes at 750 μg/mL and deposited samples immediately onto glow-discharged 1.2/1.3 quantifoil 400 grids (EMS). After incubating samples on grids for 7s, we blotted off excess sample and froze grids in liquid ethane using a Vitrobot (FEI). We transferred grids to liquid nitrogen for storage. We imaged cryoEM grids on a Titan Krios (FEI) with a Gatan K2 summit detector operating at 300kV. We collected 2,559 micrographs in counting mode at 29,000 nominal magnification using Leginon (Suloway *et al.*, 2005). Our total exposure time was 10.5s with a total dose of 53.1 electrons/Å_2_. After performing motion-correction and GCTF estimation, we picked particles using a difference-of-Gaussians picker (Voss *et al.*, 2009; Zhang, 2016; Zheng *et al.*, 2017). We performed initial reference-free 2D classification in Cryosparc followed by further 2D classification in Relion (Punjani *et al.*, 2017; Zivanov *et al.*, 2018). Next, we ran global 3D refinement with 3-fold symmetry expansion on the total particles after 2D cleanup. As pAb:HA complexes represented a minority of the total particles—mostly unbound HA and Fab—and were difficult to identify in 2D class averages, we turned to focus classification to guide identification, isolation, and refinement of each immune complex. In Relion, using the corresponding negative stain epitope landscape for subject 4 at day 28 as a guide, we positioned 40A sphere masks over expected pAbs and ran 3D classification without image alignment or imposed symmetry for each specificity. After identifying and classifying each unique pAb complex, we refined and post-processed each specificity with a mask around the entire immune complex. For our 4.7 Å resolution map of the stem-specific immune complex, we generated a predicted model of mAb 1C01 from subject 4 at day 7 using ROSIE (The Rosetta Online Server that Includes Everyone) and used Coot to minimally adjust the flexible CDR H3 loop into density (Emsley and Cowtan, 2004; Lyskov *et al.*, 2013; Weitzner *et al.*, 2017). As our resolution was not high enough, we did not further refine a model of 1C01 into the stem immune complex map.

## QUANTIFICATION AND STATISTICAL ANALYSIS

We conducted statistical analyses in GraphPad Prism.

## ADDITIONAL RESOURCES

More information about this clinical trial is located at clinicaltrials.gov under the identifier NCT01910519.

---

**Supplemental Figure 1.**
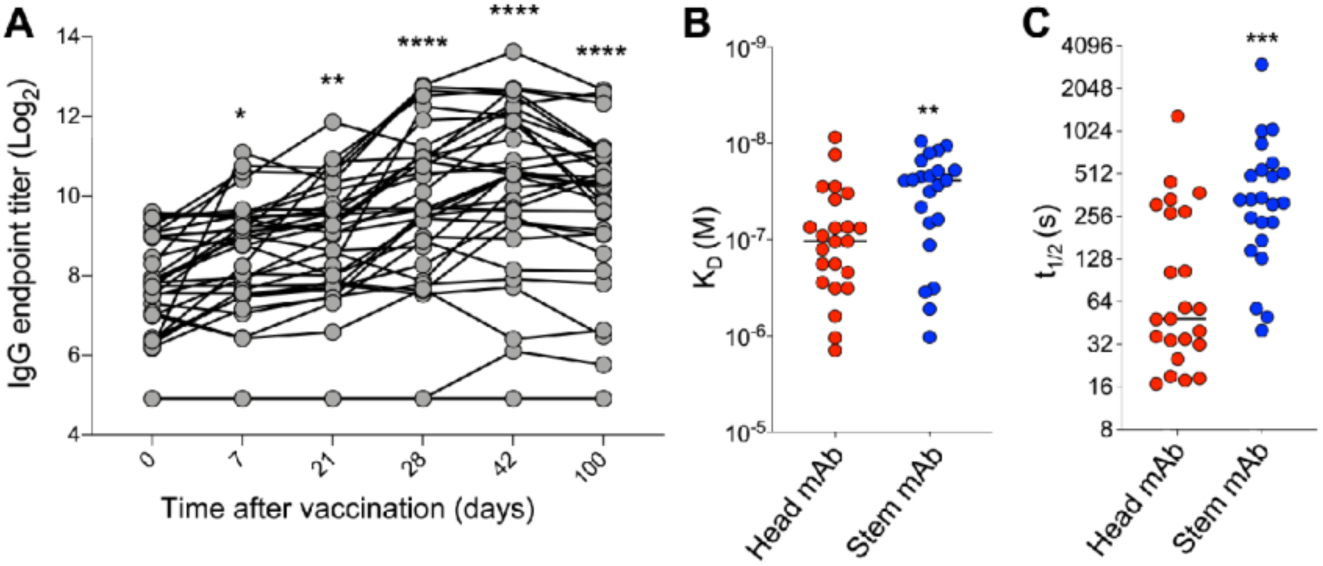
Related to Figure 1. Affinity assessment of mAbs from subjects 4, 28, 36, and 43. (A) ELISA binding titers of serum IgG to recombinant H5 HA over the course of vaccination. (B-C) affinity measurements (B) and half-life (C) of H5 HA stem- (blue) and head-specific (red) plasmablast-derived monoclonal Fab fragments isolated at days 7 and 28 binding to recombinant H5 HA. Antibody half-lives are represented as t_1/2_ (t_1/2_ = ln2/k_d_). (D-E) Representative biolayer interferometry (BLI) binding curves of monoclonal Fab fragments 1A06 (D) and 1H01 (E) isolated from subject 43 at days 7 and 28 binding to recombinant H5 HA. p-values were determined by unpaired t tests: * p-value <0.05, ** p-value < 0.01, *** p-value < 0.001, **** p-value < 0.0001, N.S. non-significant.

**Supplemental Figure 2.**
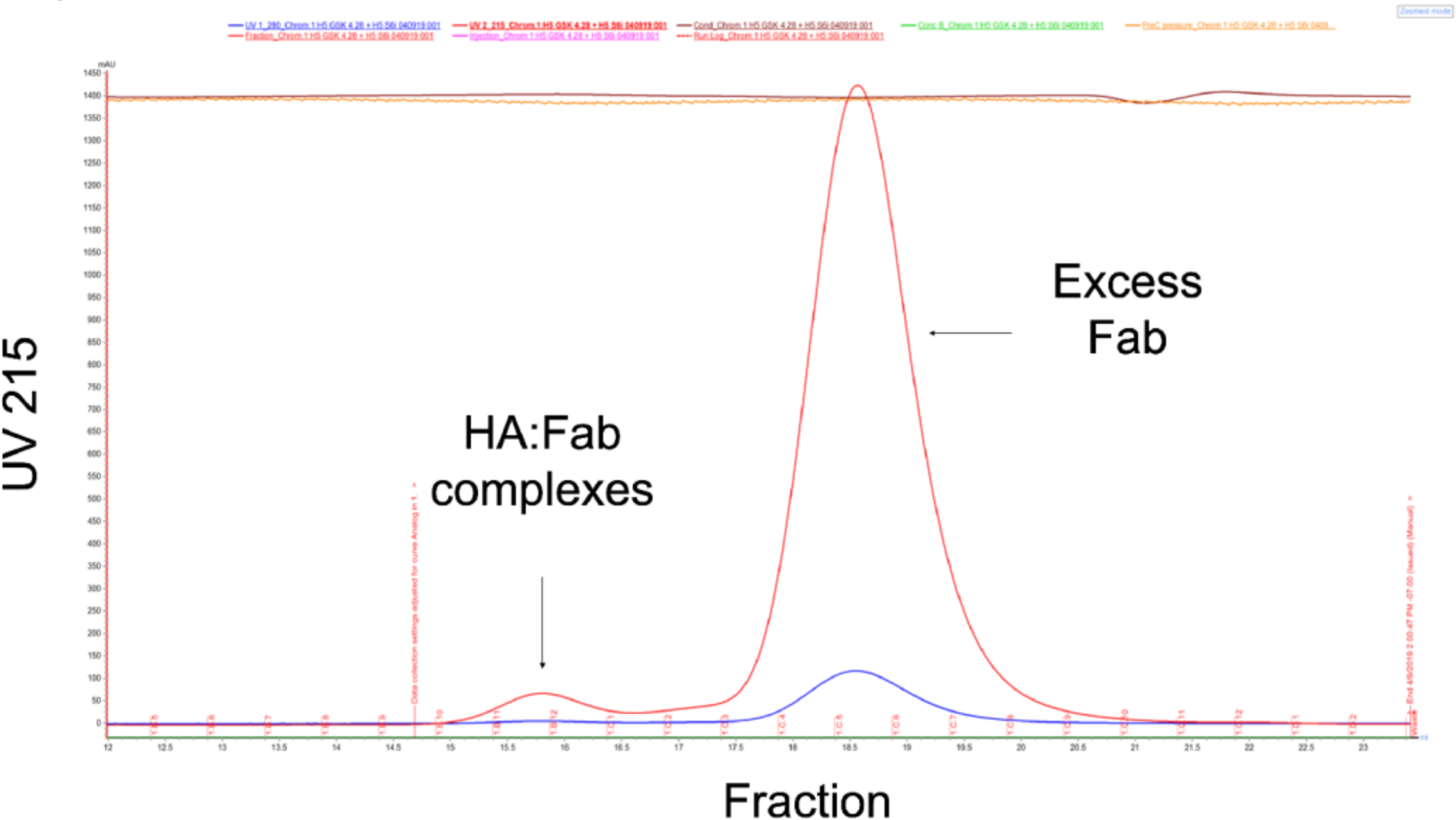
Related to Figure 2. Purification of polyclonal immune complexes. Example chromatogram from size exclusion chromatography separation of polyclonal immune complexes. Peaks corresponding to elution of immune complexes and excess Fab are labeled.

**Supplemental Figure 3.**
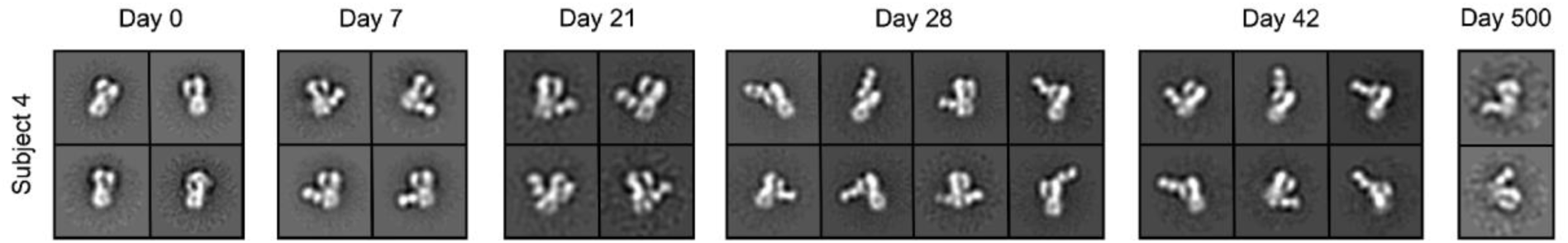
Related to Figure 3. Subject 4 2D class averages over the course of H5N1 vaccination. Example 2D class averages of immune complexes or unbound H5 HA at corresponding time points after H5N1 vaccination in subject 4.

**Supplemental Figure 4.**
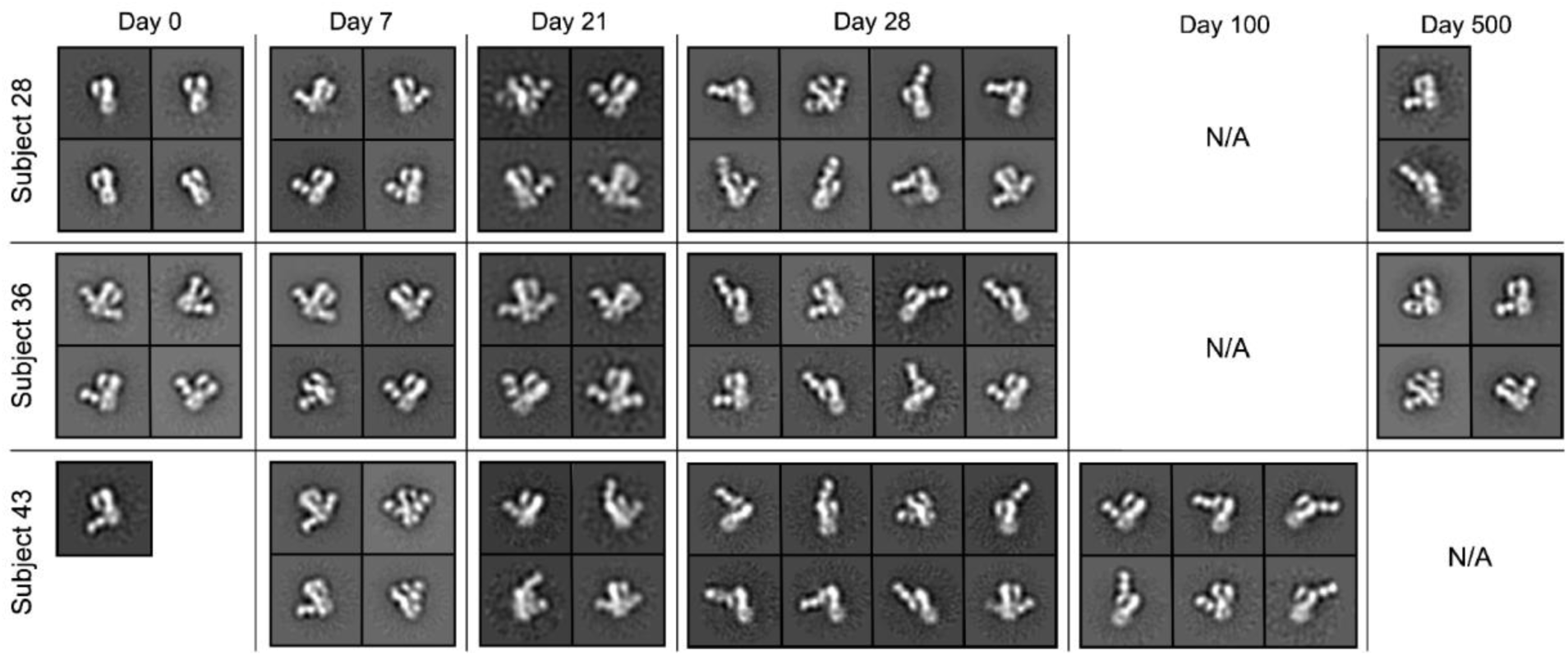
Related to Figure 4. Subjects 28, 36, and 43 2D class averages over the course of H5N1 vaccination. Example 2D class averages of immune complexes or unbound H5 HA at corresponding time points after H5N1 vaccination in subjects 28, 36, and 43.

**Supplemental Figure 5.**
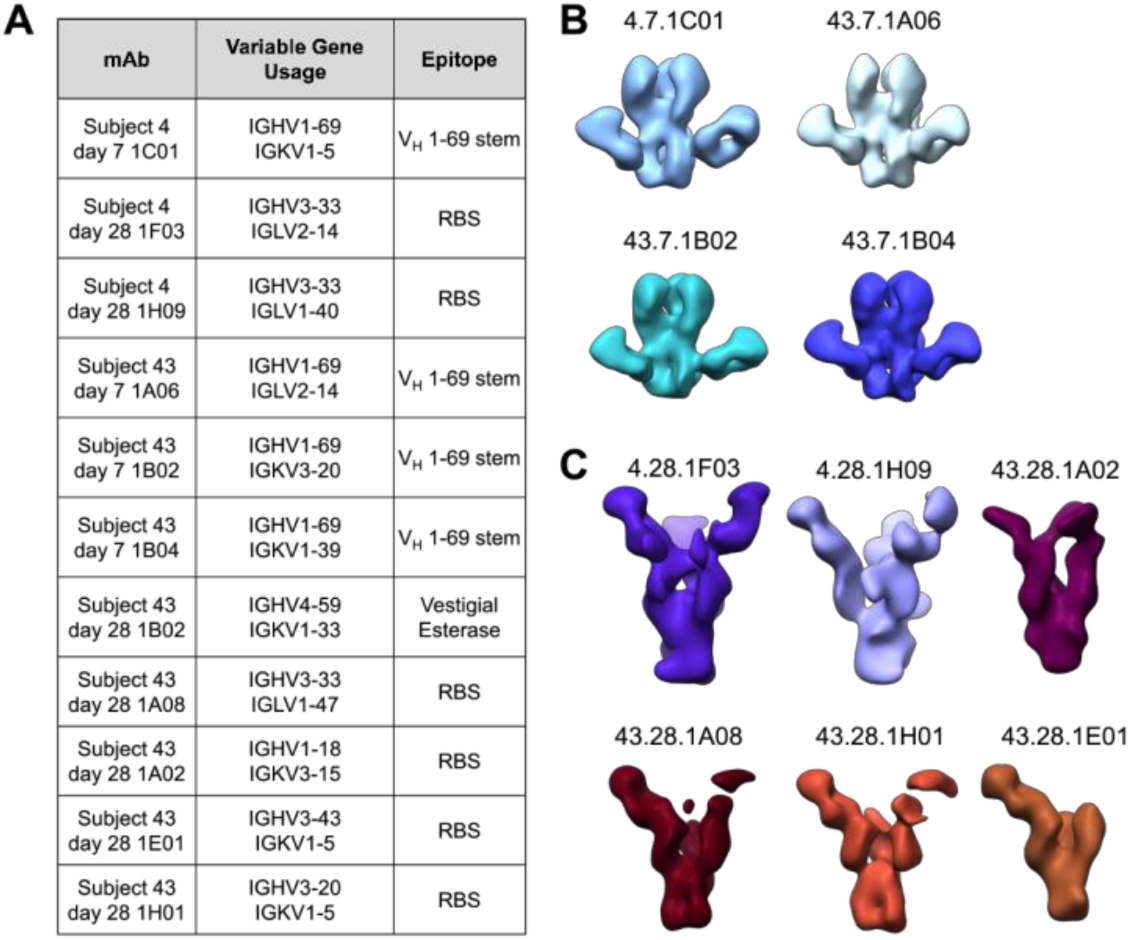
Related to Figure 5. Subject 43 mAbs target discrete regions on HA. (A) Table of mAbs isolated from plasmablasts of subjects 4 and 43. Variable gene usages and EM-mapped epitopes are listed. (B-C) 3D reconstructions of immune complexes from day 7 (B) or day 28 (C) plasmablasts with recombinant H5 HA (A/Indonesia/5/2005).

**Supplemental Figure 6.**
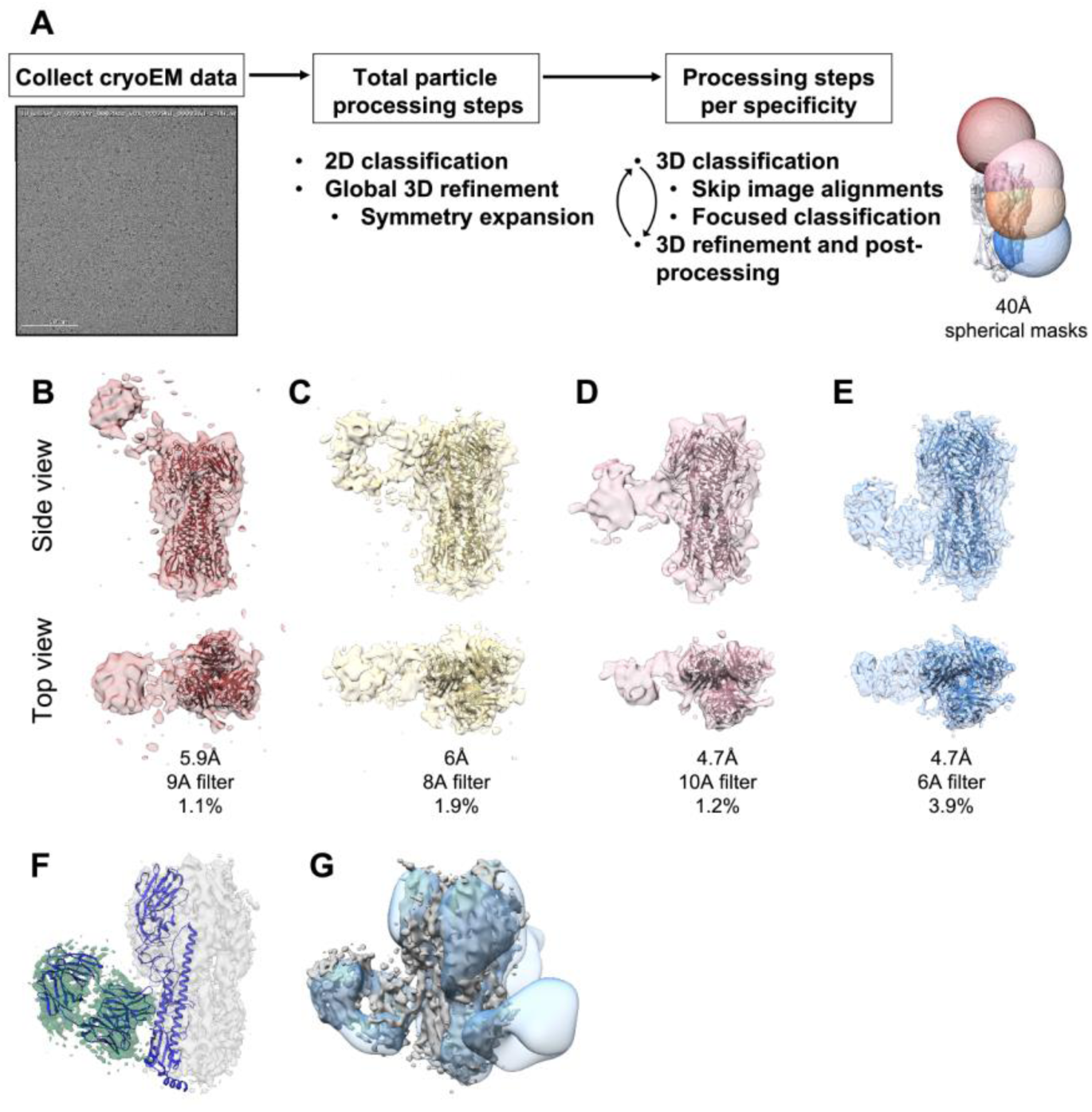
Related to Figure 6. CryoEM maps of polyclonal immune complexes. (A) Workflow of cryoEM focused classification and refinement: Briefly, after cryoEM data collection and initial processing steps, particles are classified and cleaned in 2D. All particles are masked around the HA trimer, aligned to a reference HA trimer (PDB 4K62), and C3 symmetry expanded. For focused classification, particles are classified in 3D without global image alignment using a 40 Å spherical mask around regions of anticipated pAbs. For focused refinement, 3D classes of individual immune complexes are further refined with a mask around the immune complex. Final maps of specific immune complexes are then produced from multiple iterations of focused classification and refinement. (B-E) CryoEM maps of pAbs in complex with recombinant H5 HA (A/Indonesia/5/2005). Maps were low-pass filtered to 9A for the RBS-specific (B), 8A for the lateral patch-specific (C), 10A for the vestigial esterase-specific (D), and 6A for the stem-specific (E) immune complexes. (F) CryoEM map of stem-specific immune complex in gray and green with CR9114 ribbon diagram docked into EM density (PDB 4FQI). (G) CryoEM m ap of stem-specific immune complex in grey compared with subject 4 day 7 mAb 1C01 complexed with H5 HA in blue.

**Supplemental Figure 7.**
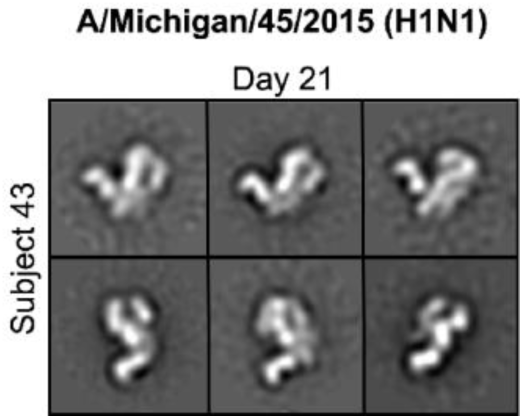
Related to Figure 7. Subject 43 2D class averages of immune complexes with heterosubtypic H1 HA. Example 2D class averages of immune complexes or unbound H1 HA (A/Michigan/45/2015) at day 21 in subject 43.

